# Polysaccharide-mediated synthesis of melanins from serotonin and other 5-hydroxy indoles

**DOI:** 10.1101/184002

**Authors:** Koen Vercruysse, Astiney Clark, Noor Alatas, Dylan Brooks, Nafisa Hamza, Margaret Whalen

## Abstract

As a continuation of our research on the melanin formation from catecholamines, we studied the polysaccharide-mediated oxidation of serotonin and other 5-hydroxy indoles into melanin-like materials. As for the catecholamines, we observed that many polysaccharides promote the oxidation of such compounds, particularly in the presence of Cu^2+^. The reactions were monitored using RP-HPLC and SEC techniques. Melanin-like materials were purified through dialysis and evaluated using UV_Vis and FT_IR spectroscopic techniques. One such material, synthesized from chondroitin sulfate type A and serotonin in the presence of Cu^2+^ was found to affect the release of IL-lβ and IL-6 cytokines from immune cells.

## 1. Introduction

In previous report, we described our observations regarding the stimulating effect many, mostly anionic, polysaccharides (PS) had on the auto-or Cu^2+^-mediated oxidation of catecholamines (CAs).[1] These reactions resulted in the formation of PS-associated, melanin (MN)- like pigments. We had hypothesized that the Cu^2+^-mediated oxidation of cationic substances like CAs was enhanced through the complexation of the cation and the substrates to the anionic PS; bringing the cationic substrates and Cu^2+^ in close proximity to each other. We have expanded these first observations by including a different cationic substrate, serotonin (5- hydroxytryptamine; 5-HT; **(1)** in Fig. 1). We evaluated the Cu^2+^-mediated oxidation of **(1)** in the presence of select PS in a similar fashion as was done for the CA substrates. In addition, we included 5-hydroxy-indole, **(2)** in Fig. 1, as a neutral substrate and 5-hydroxy-indole-3-acetic acid, **(3)** in Fig. 1, as an anionic substrate in our experiments.

**Fig. 1:**
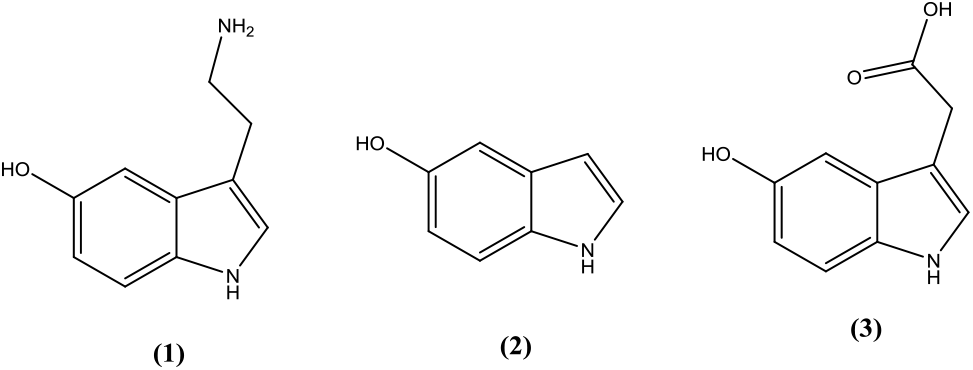
Chemical structures of the compounds used in this study. **(1)** = serotonin, **(2)** = 5-hydroxyindole and **(3)** = 5-hydroxyindole-3-acetic acid.

**(1)** is a powerful biochemical with a multitude of physiological functions as testified by the broad diversity of receptors for which this compound is a ligand.[2] Of relevance for this report is the possible involvement of **(1)** in the formation of the neuromelanins (NMs). NMs are pigmented substances found in select portions of the brain and are thought to be related to the peripheral MNs found in skin, hair, inner ear or iris.[3] The largest or densest accumulation of NMs has been found in the substantia nigra (SN) and locus coeruleus (LC) portions of the brainstem.[4] NM-containing neurons are associated with neurodegenerative diseases like Parkinson’s (PD) or Alzheimer’s disease.[3, 4] PD is characterized by the demise of melanised, dopamine-secreting neurons in select parts of the brain, e.g., the SN. The NM pigment found in these select parts of the brain is derived from CAs like dopamine or norepinephrine and its functions are unknown. Suggestions have been made that NM acts as an antioxidant or functions as storage for metal ions. However, melanised neurons appear to be more vulnerable to demise than lesser melanised neurons.[5] In addition to the CAs mentioned above, **(1)** has been investigated as a potential precursor for the formation of NMs.[6-8] Bertazzo et al. studied melanogenesis using **(1)** and related molecules as substrates for tyrosinase and peroxidase enzymes.[6] In these experiments, oligomerization of **(1)** by peroxidase was observed. Miller et al. explored the potential neurotoxicity of MNs derived from **(1)**.[9] In their studies MNs derived from **(1)** interacted with unilamellar vesicles made from the phospholipid 1,2-dioleoyl-*sn*-glycerol-3-phosphocholine, suggesting that such MNs could disrupt cellular membranes.

The reaction between Cu^2+^ and indole compounds has been studied as far back as 1960; including the observations that darkly pigmented materials can be generated.[10, 11] Using ^1^H NMR and mass spectroscopy, Jones et al. studied the reaction between Cu^2+^ and **(1)**.[12] They reported on the formation of a dimeric species, but no other oligomeric or polymeric species, and reported on the formation of an insoluble, dark brown material. They also observed that incubating rat pheochromacytoma cells with **(1)** and Cu^2+^ resulted in a reduced viability of such cells.

With this report on **(1)** and related compounds, we would like to expand our hypothesis that many PS can promote the formation of MN-like pigments as was observed for CAs and suggest the possibility that the formation and disposition of such pigments in a tissue may dependent on the PS present in the environment.

## 2. Materials and Methods

### 2.1 Materials

Chondroitin sulfate type A (sodium salt from bovine trachea; 70% with counterbalance chondroitin sulfate type C), chondroitin sulfate type C (sodium salt from shark cartilage; 90% with counterbalance chondroitin sulfate type A), alginic acid, sodium salt (Algin^®^, sodium alginate), ι-carrageenan (commercial grade, type II), fucoidan (from *Fucus vesiculosus*), serotonin.HCl, 5- hydroxyindole and 5-hydroxyindole-3-acetic acid were obtained from Sigma-Aldrich (Milwaukee, WI). CuCl_2_.2H_2_O was obtained from Fisher Scientific (Suwanee, GA). All other reagents were of analytical grade.

### 2.2 Stock solutions and reaction mixtures

Stock solutions of CuCl_2_.2H_2_O were prepared in water in advance. Stock solutions of all PS were prepared in water one day prior to the start of the experiment. The pH of these PS solutions was measured and ranged between 6.4 and 7.5. Stock solutions of serotonin.HCl were prepared in water just prior to the start of the experiments. Stock solutions of 5-hydroxy indole and 5-hydroxy indole-3-acetic acid were prepared in methanol:water (1:9 v/v) just prior to the start of the experiments.

### 2.3 UV/Vis spectroscopy

UV/Vis spectroscopy was performed in microwells of a 96-well microplate using the SynergyHT microplate reader from Biotek (Winooski, VT). 200 μL aliquots were used and 200 μL water as the blank. All measurements were performed at room temperature (RT).

### 2.4 RP-HPLC analyses

RP-HPLC analyses were performed on a UFLC chromatography system equipped with dual LC-6AD solvent delivery pumps and SPD-M20A diode array detector from Shimadzu, USA (Columbia, MD). Analyses were performed on a BDS Hypersil C_8_ column (125X4.6 mm) obtained from Fisher Scientific (Suwanee, GA). Analyses were performed in isocratic fashion using a mixture of water:methanol:acetic acid (90:10:0.05% v/v) as solvent. The sample volume was 20 μL. Samples were diluted to an approximate final concentration of 0.1mM for **(1)**, **(2)** or **(3)** and centrifuged prior to analysis.

### 2.5 Size exclusion chromatography (SEC)

SEC analyses were performed on a Breeze 2 HPLC system equipped with two 1500 series HPLC pumps and a model 2998 Photodiode array detector from Waters, Co (Milford, MA). Analyses were performed using an Ultrahydrogel 500 column (300 X 7.8 mm) obtained from Waters, Co (Milford, MA) and in isocratic fashion using a mixture of 25mM Na acetate:methanol:acetic acid (90:10:0.05% v/v) as solvent. Samples were diluted with SEC solvent to an approximate final concentration of 0.1mM for **(1)**, **(2)** or **(3)** and centrifuged prior to analysis, centrifuged. The sample volume was 20 μL.

### 2.6 Dialysis and freeze drying

Select samples were dialyzed using Spectrum Spectra/Por RC dialysis membranes with molecular-weight-cut-off of 3.5kDa obtained from Fisher Scientific (Suwanee, GA). Select dialyzed materials were frozen overnight and dried using a Labconco FreeZone Plus 4.5L benchtop freeze-dry system obtained from Fisher Scientific (Suwanee, GA). All dried materials were kept in a freezer till further analysis.

### 2.7 FT-IR spectroscopy

FT-IR spectroscopic scans were made using the NicoletiS10 instrument equipped with the SmartiTR Basic accessory from ThermoScientific (Waltham, MA). Scans were taken with a resolution of 4 cm^-1^ between 650 and 4,000 cm^-1^ at room temperature using a KBr beam splitter and DTGS KBr detector. Each spectrum represents the accumulation of 24scans.

### 2.8 Atomic absorption spectroscopy (AAS)

AAS measurements were made using the AAnalyst 300 instrument from Perkin Elmer (Waltham, MA). Standard solutions containing known concentrations of CuCl_2_ were prepared in water and used to calibrate the instrument for the analyses. Stock solution of pigment material was prepared in water and analyzed as such.

### 2.9 Preparation of monocyte-depleted (MD) peripheral *blood mononuclear cells (PBMCs)*

PBMCs were isolated from Leukocyte filters (PALL-RCPL or RC2D) obtained from the Red Cross Blood Bank Facility (Nashville, TN) as described elsewhere.[13] Leukocytes were retrieved from the filters by back-flushing them with an elution medium (sterile PBS containing 5 mM disodium EDTA and 2.5% [w/v] sucrose) and collecting the eluent. The eluent was layered onto Ficoll-Hypaque (1.077g/mL) and centrifuged at 1,200g for 50 min. Granulocytes and red cells pelleted at the bottom of the tube while the PBMCs floated on the Ficoll-Hypaque. Mononuclear cells were collected and washed with PBS (500g, 10min). Following washing, the cells were layered on bovine calf serum for platelet removal. The cells were then suspended in RPMI-1640 complete medium which consisted of RPMI-1640 supplemented with 10% heat-inactivated BCS, 2 mM *L*-glutamine and 50 U penicillin G with 50 μg streptomycin/mL. This preparation constituted PBMCs. Monocyte-depleted PBMCs (10-20% CD16^+^, 10-20 % CD56^+^, 70-80% CD3^+^, 3-5% CD19^+^, 2-20% CD14^+^) were prepared by incubating the cells in glass Petri dishes (150 X 15 mm) at 37 °C and air/CO_2_, 19:1 for 1 h. This cell preparation is referred to as MD-PBMCs.

### 2.10 PBMC studies

Test compounds were dissolved in water one day prior to the start of the experiment. For the IL-1β release experiment, 25 μL of the test compounds (and the appropriate controls) were added to 500 μL of MD-PBMC cell suspension. For the IL-6 release, 5 μL of the test compounds (and the appropriate controls) were added to 1,0 μL of MD-PBMC cell suspension. The concentration of the cells was 750,000 in 500 μL of cell culture media. After addition of the compounds, the mixtures were incubated at 37°C in an atmosphere containing 5% CO_2_ for 24 h after which cell viability was assessed and supernatants were collected and stored at - 80°C until assay.

### 2.11 Cell Viability

Cell viability was assessed at the end of each exposure period. Viability was determined using the trypan blue exclusion method. Briefly, cells were mixed with trypan blue and counted using a hemocytometer. The total number of cells and the total number of live cells were determined for both control and treated cells to determine the percent viable cells

### 2.12 IL-1ß or IL-6 secretion assay

IL-1β and IL-6 levels in isolated samples were assessed using the OptEIA™ enzyme-linked immuno-sorbent assay (ELISA) for human IL-1β kit and human IL-6 respectively (BD Biosciences, San Diego, CA). In brief, capture antibody diluted in coating buffer was applied to wells of 96-well flat-bottom micro-plates specifically designed for ELISA (Fisher Scientific). The plates were incubated overnight at 4°C, and then excess capture antibody was removed by washing the plate three times with wash buffer (PBS containing 0.05% [v/v] Tween-20 (PBST)). The wells were then treated with blocking buffer to prevent non-specific binding and the plate was sealed and incubated at room temperature for 1 hr. Blocking buffer was removed with three washes of PBST, and cell supernatants and IL-1β or IL-6 standards were added to dedicated wells; the plate was re-sealed and incubated at room temperature for 2 hr. The plate was then washed five times with PBST and this was followed by incubation with IL-1β or IL-6 detection antibody which was subsequently conjugated with horseradish peroxidase. The plate was then washed seven times and a substrate solution was added for 30 min at room temperature to produce a colored product. The incubation with the substrate was ended by addition of acid and the absorbance was measured at 450 nm on a Thermo Labsystems Multiskan MCC/340 plate reader (Fisher Scientific).

## 3. Results

### 3.1 Preliminary studies using RP-HPLC

When aqueous solutions of **(1)**, **(2)** or **(3)** (concentrations about 1mM) are kept at RT or 37°C, with or without the presence of Cu^2+^ (concentrations below 0.1mM), then little change in color of these solutions was observed. However, when such solutions contained select PS, then slowly, typically after overnight reactions, dark brown to black colors appeared in some of these mixtures. Such observations hinted to the possibility that select PS, e.g., chondroitin sulfate type A (CS A) or fucoidan (FUCO) could promote the Cu^2+^-mediated oxidation of **(1)** and related compounds to MN-like pigments, similar to what was observed for reactions between PS and CAs.[1] Aliquots from such reaction mixtures could be analyzed using RP-HPLC or SEC. Typical RP-HPLC profiles are shown in Fig. 2.

**Fig.2.**
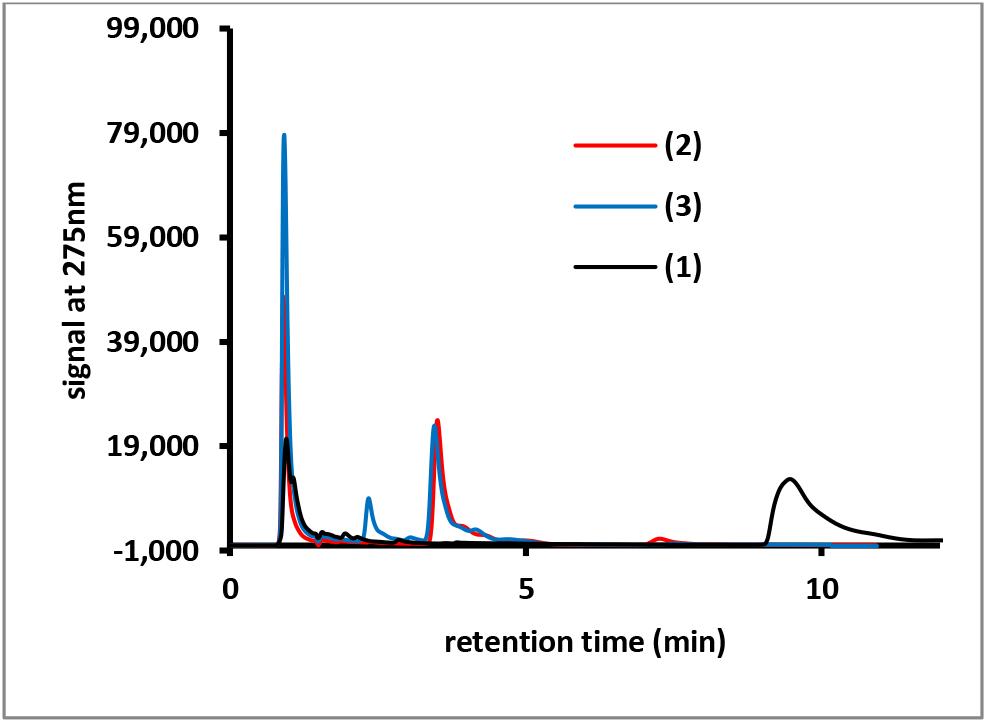
RP-HPLC profiles of aliquots of reaction mixtures containing CS A, Cu^2+^ and **(1)** (black line), **(2)** (red line or **(3)** (blue line). The reaction mixtures contained 1mM **(1)**, **(2)** or **(3)**, 8 mg/mL CS A and 0.1mM Cu^2+^ and were kept at 37°C overnight.

In such analyses **(1)** had a retention time of about 9.5 minutes, while **(2)** and **(3)** had retention times of about 3.75 minutes. RP-HPLC profiles of reaction mixtures containing PS would show peaks with retention times of about 1 minute that are associated with the PS material. RP-HPLC profiles from reaction mixtures involving **(2)** or **(3)** contained peaks of unknown identity with retention times of about 7.5 or 2.5 minutes respectively. Mixtures containing 1mM **(1)**, 2.5mg/mL CS A and between 0 and 1.3 mM Cu^2+^ were incubated overnight at 37°C. Aliquots of the reaction mixtures were diluted and analyzed using RP-HPLC and the AUC for the signal at 275nm for the peak corresponding to **(1)** was calculated. In addition, the absorbance at 400nm of the undiluted reaction mixtures was measured in a microplate. Fig.3 presents the results thus obtained.

**Fig. 3:**
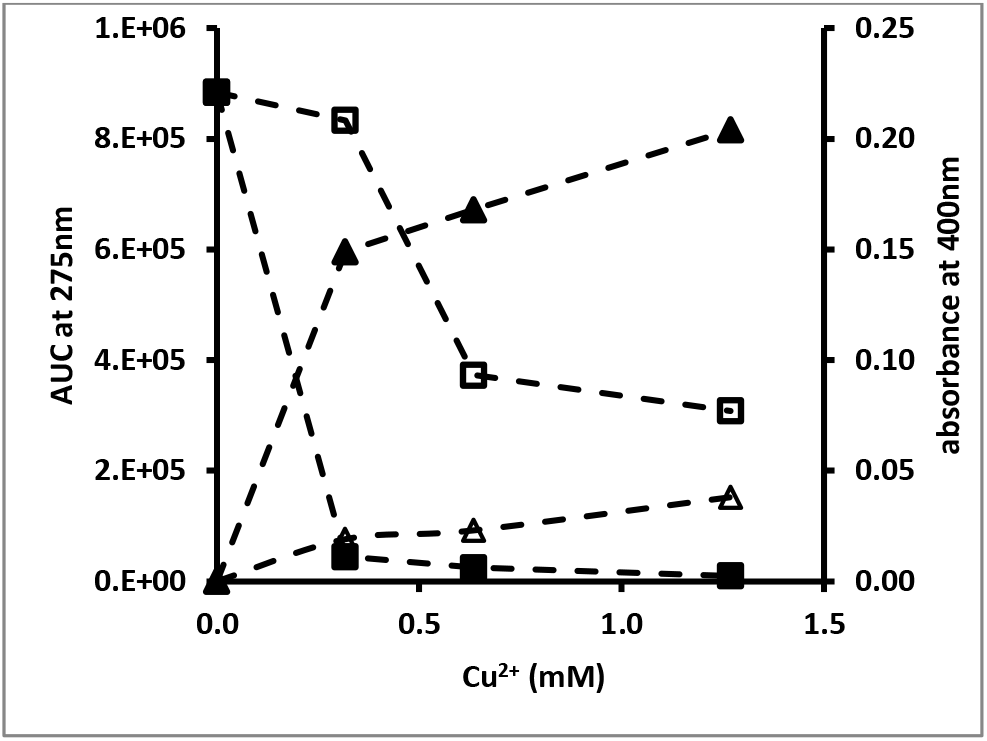
Effect of Cu^2+^ concentration on reactions containing **(1)**, CS A and Cu^2+^. AUC corresponding to **(1)** for reactions with (closed squares) or without (open squares) CS A and absorbance at 400nm of the reaction mixtures with (closed triangles) or without (open triangles) CS A. The reaction mixtures contained 1mM **(1)**, 2.5mg/mL CS A and between 0 and 1.3 mM Cu^2+^ and were incubated overnight at 37°C.

In the absence of any CS A, a decline (up to 60%) in AUC of the peak corresponding with **(1)** and an increase in absorbance at 400nm could be observed with increasing Cu^2+^ concentrations. In the presence of CS A a near total disappearance of **(1)** from the reaction mixtures was observed in the presence of the highest concentration of Cu^2+^ tested. This was associated with the presence of a much darker color of the reaction mixtures as judged from visual observations and by measuring the absorbance of the mixtures at 400nm. Clearly, in the presence of CS A, a much stronger reactivity between **(1)** and Cu^2+^ had occurred. A similar pattern of results was obtained for reactions in the presence of chondroitin sulfate type C (CS C), carrageenan (CARRA), alginate (ALG) or fucoidan (FUCO) (results not shown). In addition, CARRA and FUCO appeared to promote color formation from **(1)** in the absence of any added Cu^2+^ (results not shown).

The effect of Cu^2+^ concentration on the reaction with **(2)** or **(3)** in the presence of CS A was briefly investigated. Higher concentrations of CS A were used in these experiments as visual observations made during preliminary experiments had indicated that **(2)** and **(3)** reacted less compared to **(1)**. Mixtures containing 1mM **(2)** or **(3)**, between 0 and 0.17mM Cu^2+^ and 0 or 8 mg/mL CS A were incubated overnight at 37°C. Aliquots from the reaction mixtures were diluted and analyzed using RP-HPLC. The AUC (signal at 275nm) for the peaks corresponding to **(2)** or **(3)** and the unknown reaction product with retention time of about 7.5 or 2.5 minutes (see Fig.2) were determined. Fig.4, panel A, illustrates the decline in AUC of the peak corresponding to **(2)** and the increase in the AUC of the peak corresponding to the reaction product formed as a function of the Cu^2+^ concentration present in the mixtures. Fig. 4, panel B, illustrates similar results for the reactions involving **(3)**.

**Fig. 4:**
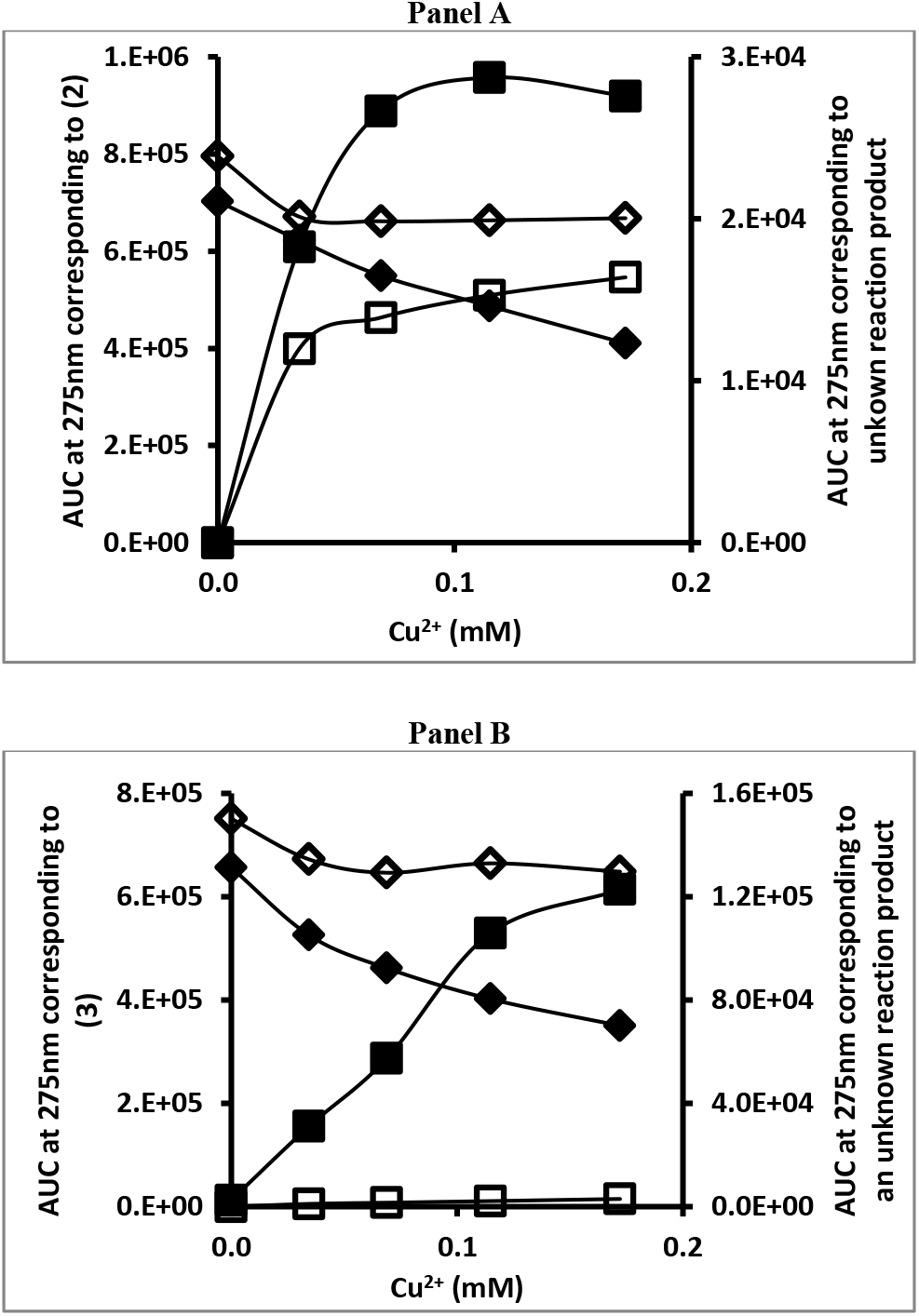
Effect of Cu^2+^ concentration on the reaction between **(2)** (panel A) or **(3)** (panel B), CS A and Cu^2+^. AUC corresponding to **(2)** or **(3)** (diamonds) for reactions with (closed symbols) or without (open symbols) CS A and AUC corresponding to an unknown reaction product (squares) as a function of Cu^2+^ concentration present. The reaction mixtures contained 1mM **(2)** or **(3)**, 0 or 8mg/mL CS A and between 0 and 0.17 mM Cu^2+^ and were incubated overnight at 37°C.

The results indicate that in the presence of CS A, there was a Cu-dependent decline in the AUC of the peak corresponding to **(2)** or **(3)**. However, there was no near-total disappearance of the substrate as was observed for the experiments involving **(1)**. In the absence of CS A, a modest decline in the AUC corresponding to **(2)** or **(3)** was observed as a function of the Cu^2+^ concentration. For the experiments involving **(2)**, there was a Cu^2+^-dependent increase in AUC of the peak corresponding to the reaction product formed, but this increase was much higher in the presence of CS A than in the absence of this PS. For the experiments involving **(3)**, there was a Cu^2+^-dependent increase in AUC of the peak corresponding to the reaction product formed in the presence of CS A, but hardly any such reaction product was observed in the absence of the PS.

### 3.2 Preliminary studies using SEC

As discussed for the experiments involving CAs, SEC is a versatile technique to monitor: (a) the presence and disappearance of the substrate of the reaction, (b) the characteristics of the PS-associated materials, (c) the potential presence of nanoparticles (NPs) and (d) the presence of other reaction products.[1] SEC analyses for reaction mixtures containing **(2)** were not routinely performed as this compound appeared to exhibit an extensive retention on the SEC column used and its retention time for these analyses could not be determined. Fig. 5 shows typical SEC profiles for reactions involving **(1)** or **(3)** in the presence of CS A and Cu^2+^.

**Fig. 5:**
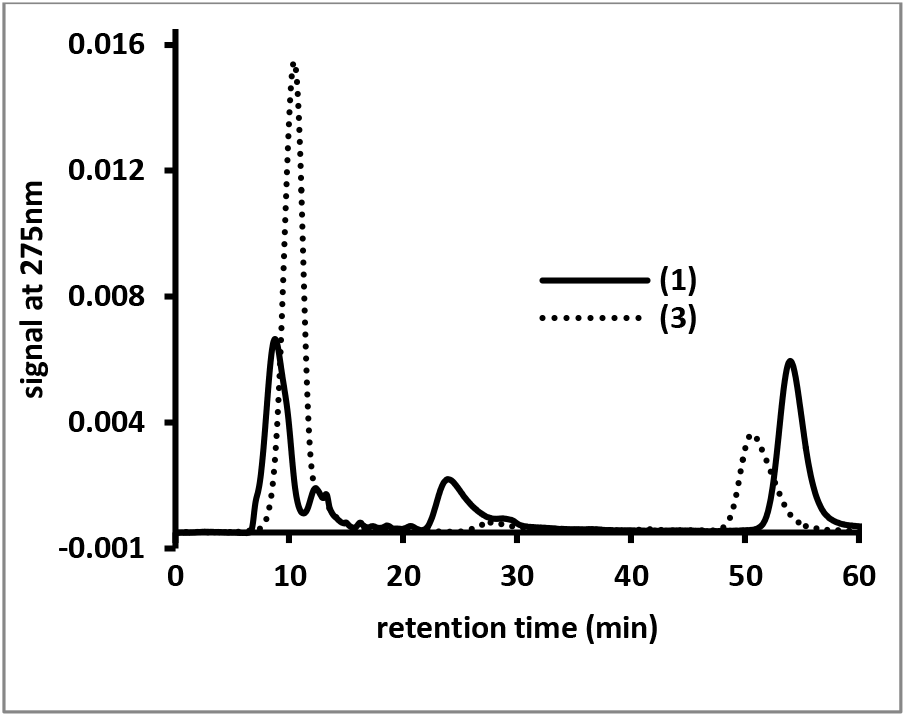
SEC profiles of aliquots of reaction mixtures containing CS A, Cu^2+^ and **(1)** (solid line) or **(3)** (dotted line). The reaction mixtures contained 1mM **(1)** or **(3)**, 8 mg/mL CS A and 0.1mM Cu^2+^ and were kept at 37°C overnight.

The peaks with retention times of less than 15 minutes correspond to high molecular mass materials, e.g., PS or PS/pigment complexes. The peaks with retention times of about 51 or 57 minutes correspond to **(3)** or **(1)** respectively. The peaks with retention times of about 25 or 28 minutes correspond to unidentified reaction products that appeared during the reactions involving **(1)** or **(3)** respectively. An unknown reaction product was observed in the RP-HPLC profiles for reaction mixtures involving **(3)** (see Figure 3), but not for the reaction mixtures involving **(1)**.

A kinetic study was set up involving reactions containing 2mM **(1)**, 4 mg/mL CS A or FUCO and 0 or 0.05mM Cu^2+^ incubated at 37°C. Over a period of multiple days, aliquots from these reaction mixtures were diluted, centrifuged and analyzed using SEC. Fig. 6 illustrates the AUC (signal at 275nm) of the peak corresponding to **(1)** as a function of the reaction time.

**Fig.6:**
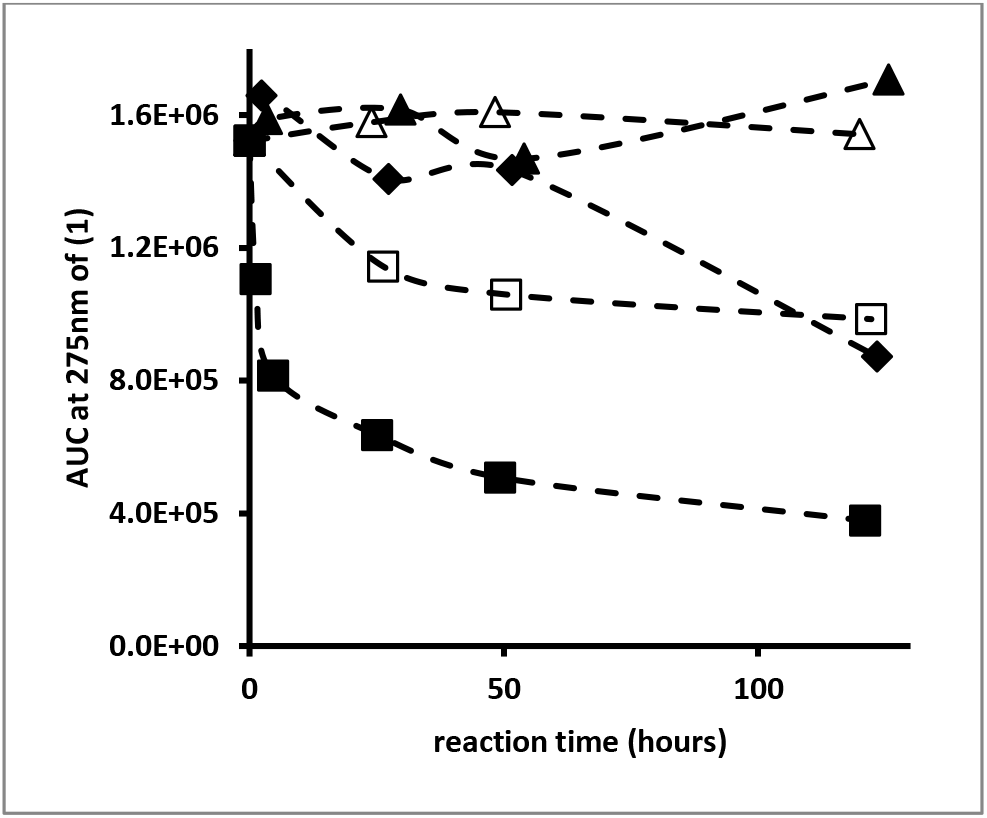
Kinetic profile of the AUC (signal at 275nm) corresponding to **(1)** as determined by SEC analysis. Reaction mixtures contained 2mM **(1)** and 0 (open symbols) or 0.05 (closed symbols) mM Cu^2+^ and 0 mg/mL PS (triangles), 4 mg/mL chondroitin sulfate A (diamonds) or 4mg/mL fucoidan (squares).

In the absence of any PS, with or without Cu^2+^, no significant decline in AUC related to **(1)** could be observed. The reaction in the presence of CS A and Cu^2+^ resulted in a decline of about 40% in the AUC associated with **(1)** at the end of the experiment. In the presence of CS A, without any added Cu^2+^, no significant decline in AUC associated with **(1)** could be observed (results not included in Fig. 6). In the presence of FUCO, the AUC associated with **(1)** declined much more rapidly and by the end of the experiment had declined to about 65% of starting value in the absence of any added Cu^2+^ and about 25% of starting value in the presence of Cu^2+^. Associated with these experiments, we also monitored the AUC of the unknown reaction product with retention time of about 25 minutes and the AUC of the peak associated with PS materials. Fig. 7 illustrates these results for the reaction between **(1)** and FUCO in the presence of Cu^2+^.

**Fig. 7:**
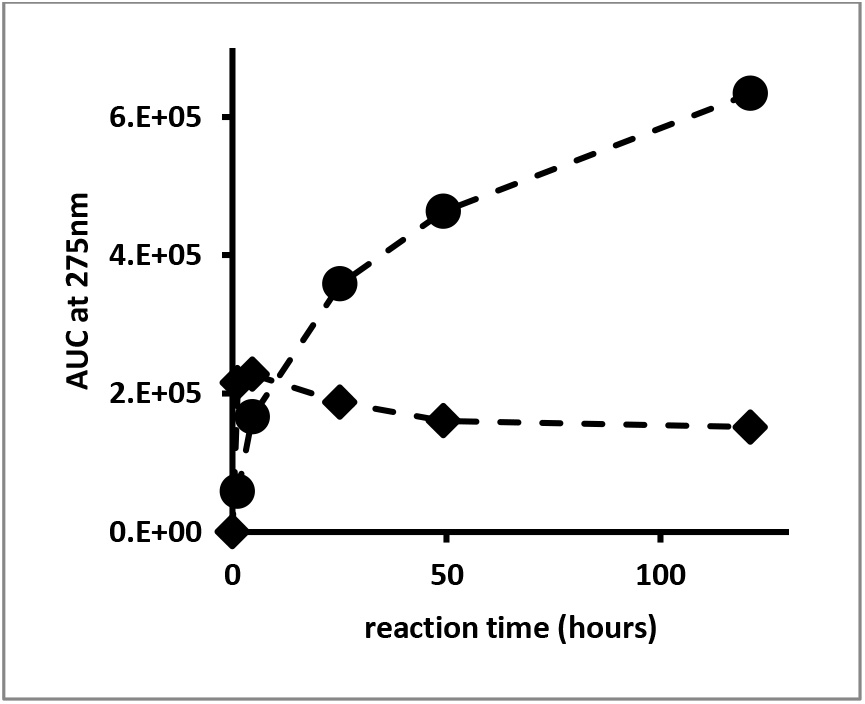
Kinetic profile of the AUC (signal at 275nm), as determined by SEC analysis, corresponding to FUCO (circles) and unknown reaction product (diamonds). The reaction mixture contained 2mM **(1)**, 0.05 mM Cu^2+^ and 4 mg/mL FUCO.

The results show a rapid increase in the AUC of the unknown reaction product, followed by a very slow decline. In addition, the UV_Vis absorbance associated with the PS materials steadily increased as a function of the reaction time. A similar profile for the changes in AUC associated with PS and reaction products had been observed for the reaction between select PS and CAs.[1]

### 3.3 Large scale reactions, isolation and characterization of pigments

A 10mL mixture containing 520 mg CS A, 1.3 mg CuCl_2_.2H_2_O (0.7mM Cu^2+^) and 30 mg **(1)** (14mM) was kept at 37°C for six days. Higher concentrations, compared to the small scale preliminary experiments, of CS A, Cu^2+^ and **(1)** were used to maximize the formation of pigment materials. Following this reaction, a photograph of the mixture was taken and is shown in Fig.8, panel A. Fig. 8, panel B, shows a photograph of the dialyzed and lyophilized material thus obtained.

**Fig. 8:**
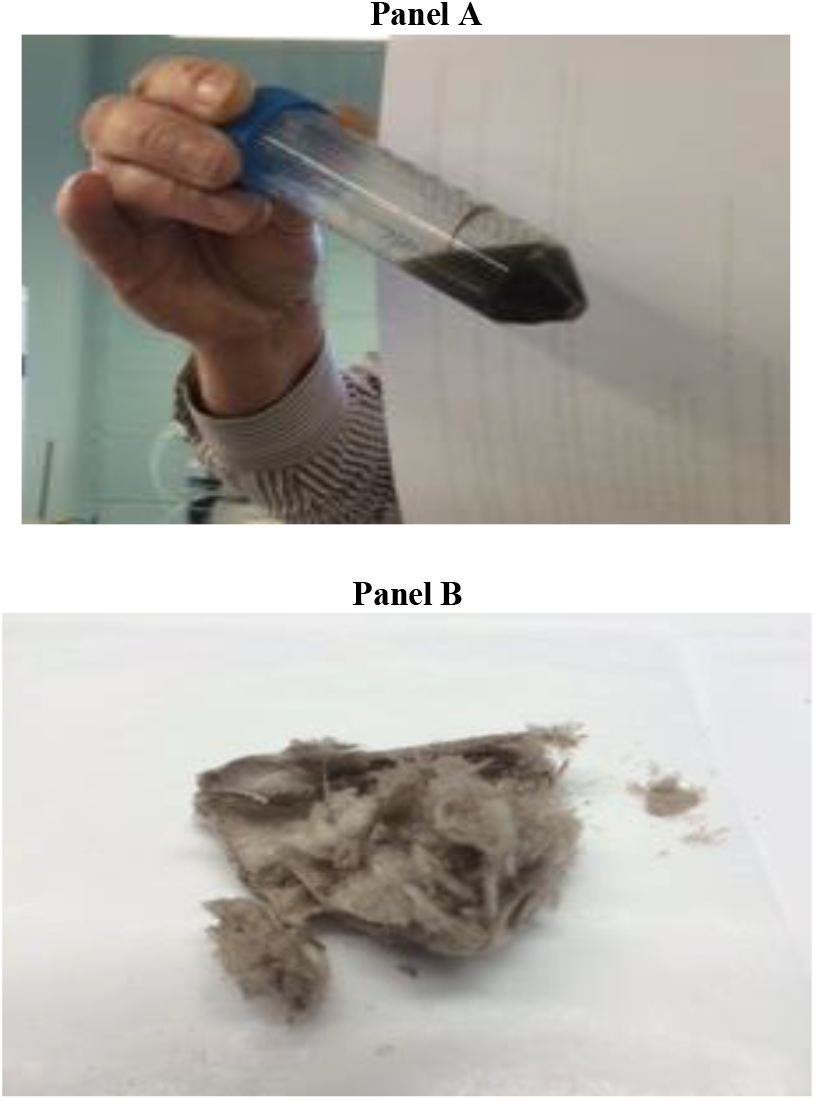
Photograph of the crude reaction mixture (Panel A) containing 14mM **(1)**, 52 mg/mL CS A and 0.7mM Cu^2+^ after six days of reaction at 37°C and of the dialyzed and dried material (Panel B).

The photograph in Fig.8, panel A, shows a darkly-colored solution and a ring of dark material adhered to the side of the plastic tube used as reaction vessel.

Fig. 9 shows the SEC profiles (signal at 275nm) of a diluted aliquot of the reaction mixture before and after exhaustive dialysis.

**Fig. 9:**
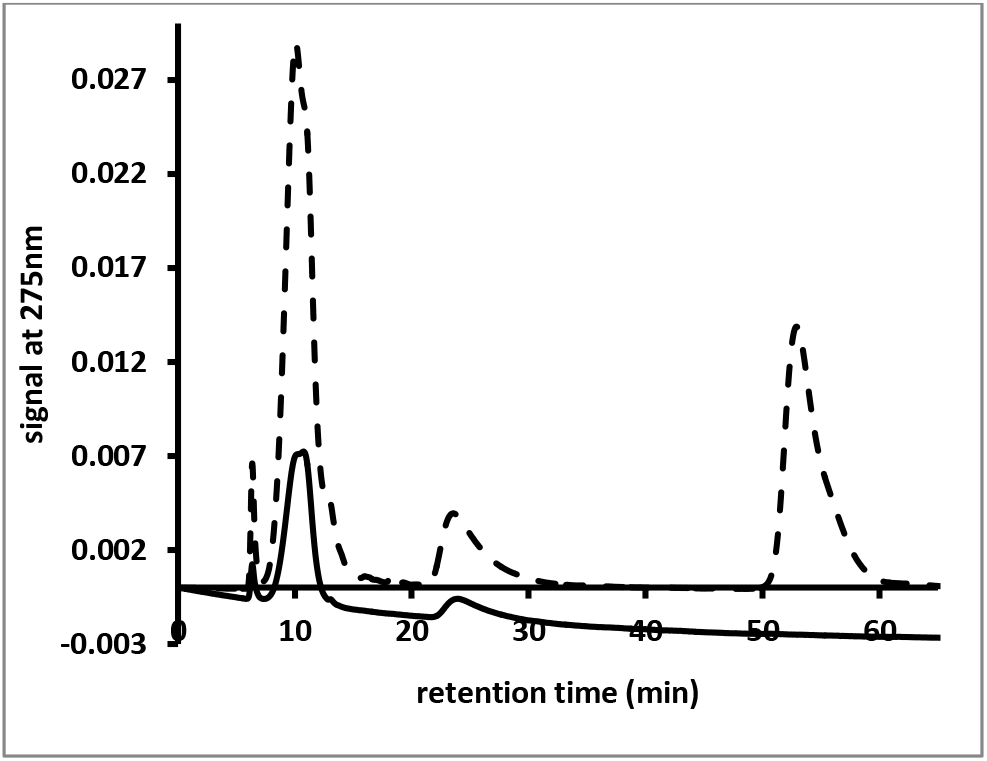
SEC profiles of diluted aliquots from the reaction mixture (dotted line) shown in Figure 8, panel A and the reaction mixture after dialysis against water (solid line).

At the end of the reaction the SEC profile shows peaks corresponding to: (a) **(1)** (retention time about 55 minutes), (b) an unknown reaction product typically observed under these reaction conditions (retention time about 25 minutes, (c) CS A (retention time about 10 minutes) and (d) a very high molecular mass material (peak retention time of about 6 minutes). Following dialysis, the mixture contains no **(1)**, but still has peaks corresponding to the three other items mentioned above. When the SEC profile of the dialyzed mixture is viewed at 400nm, then only the peaks corresponding to CS A and the very high molecular mass material are observed (results not shown). This indicates that the material with absorbance in the visible range is associated with the CS A peak and with the very high molecular mass material with peak retention time of about 6 minutes.

The dialyzed and dried material shown in Fig. 8, panel B, was redissolved in water to a concentration of 5mg/mL. AAS analysis was performed on this pigment solution and on Cu^2+^ standards with concentrations between 0 and 0.4mM. The results indicated that following exhaustive dialysis the dried pigment material contained 1.4 μg Cu^2+^/mg material.

An FT-IR spectrum of the dried pigment material was recorded and is presented in Fig. 10, panels A and B. Fig. 10, panel A, compares the FT-IR spectra of **(1)** and the pigment material described above. Fig. 10, panel B, compares the FT-IR spectra of CS A and this same pigment material.

**Fig. 10:**
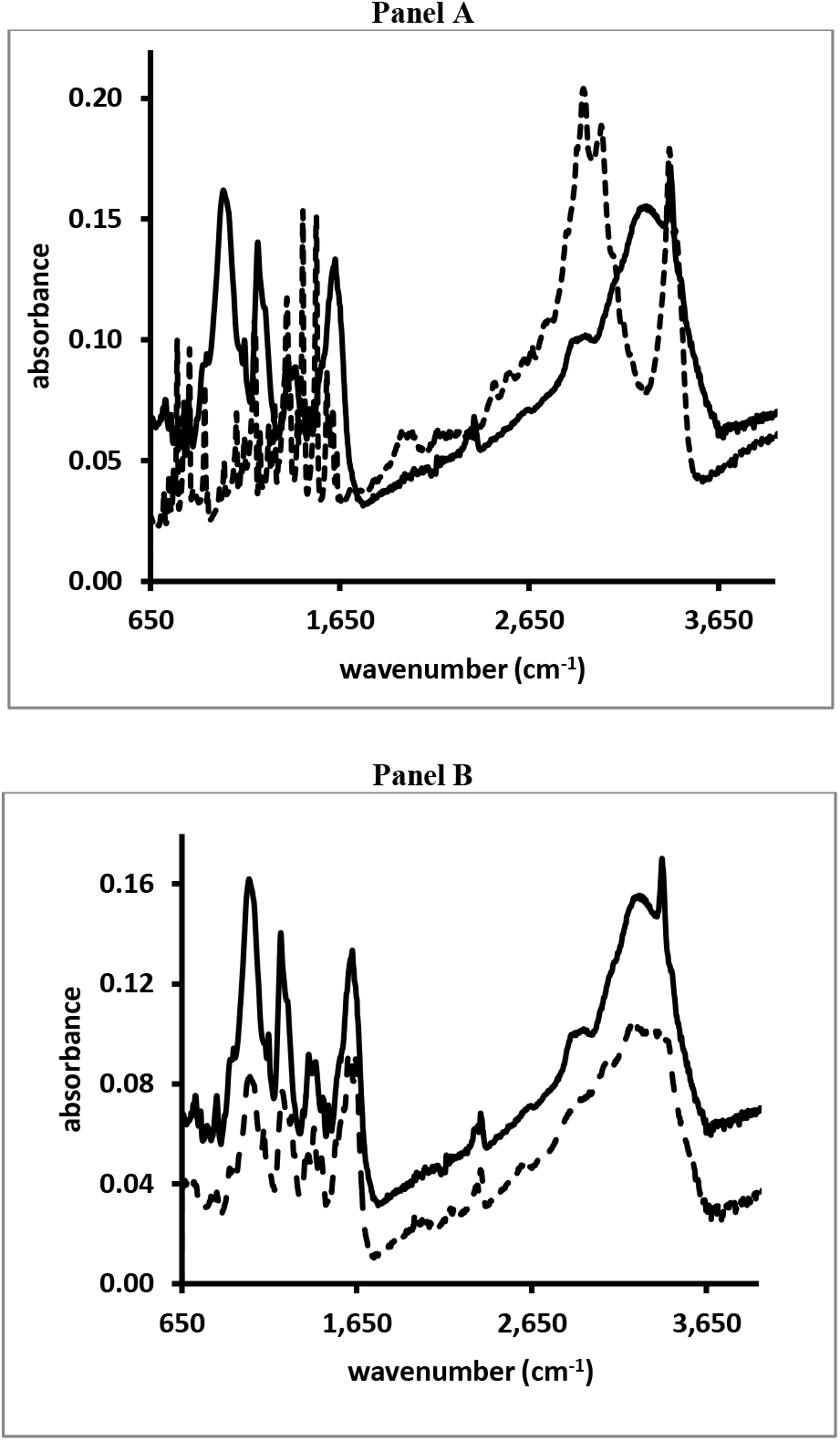
FT_IR scans of the pigment (solid line in both Panels) made from **(1)** as shown in Figure 8, panel B and **(1)** (dashed line in Panel A) or CS A (dashed line in Panel B).

The IL-1β and IL-6 release from immune cells in the presence of CS A or the pigment material discussed earlier (see Fig. 8) is shown in Fig. 11, panels A and B respectively.

**Fig. 11:**
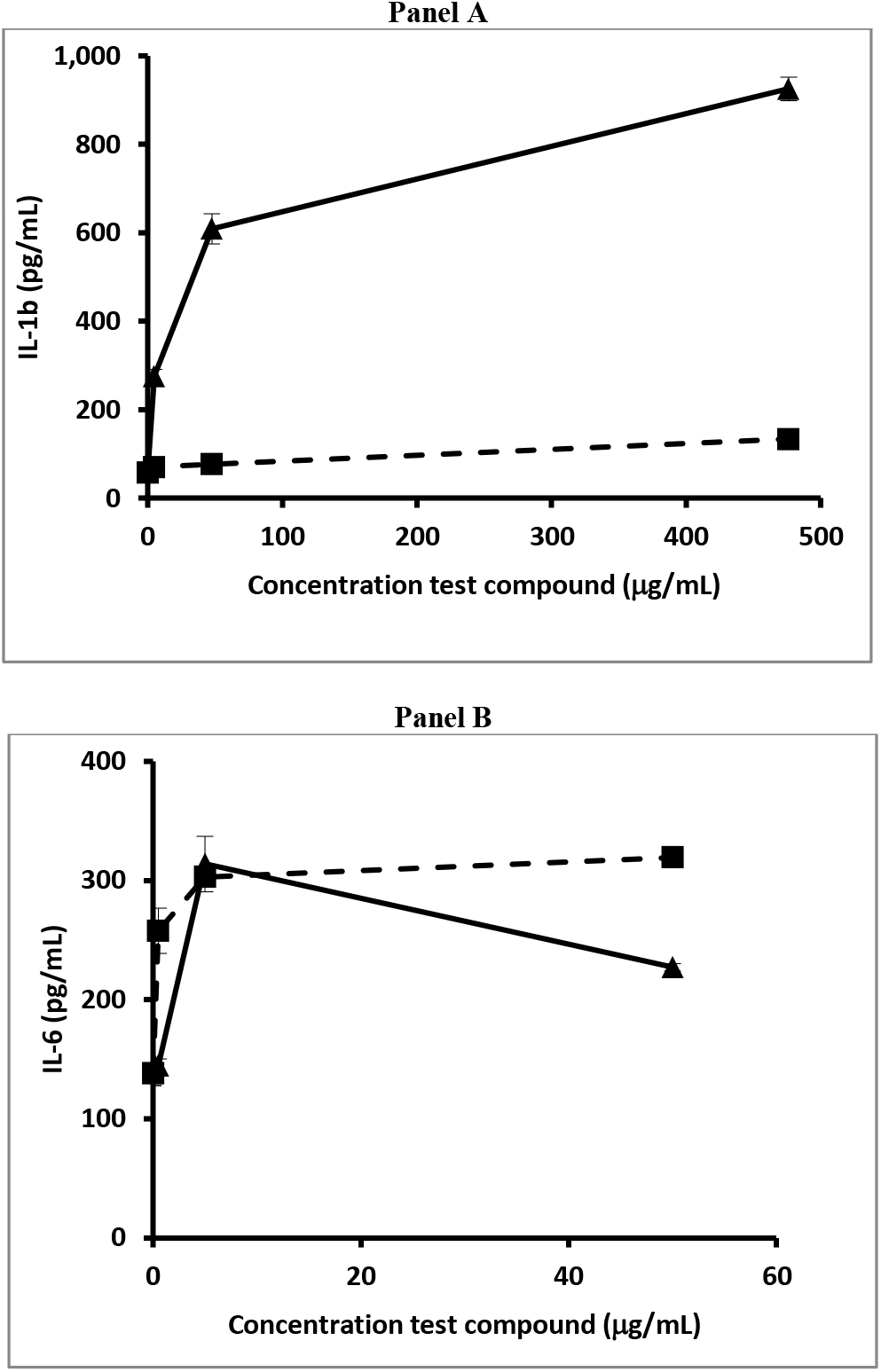
IL-1β (panel A) or IL6 (panel B) release from immune cells in the presence of CS A (solid squares) and the pigment (solid triangles) made from **(1)** and CS A in the presence of Cu^2+^ as shown in Fig. 8.

CS A and the dried pigment were dissolved in water at a concentration of 10 mg/mL and a ten-and hundred-fold dilution of both stock solutions were prepared for this experiment. The results show that, while CS A had a minimal effect on the IL-1β release from the immune cells, the pigment material induced the release of significant amounts of IL-1β. In addition, both CS A and the pigment material did not affect the viability of the immune cells. In the case of Il-6 release, the effect of the pigment material appeared to deviate from the effect of CS A when the pigment was tested at a higher concentration.

Large scale reactions were set up between **(2)** or **(3)** and select PS or sodium acetate in the absence of any added cation. Between 30 and 35 mg of **(2)** or **(3)** was placed into the well of a cell culture dish and 10mL water was added. To these, about 100 mg of CS A, CS C, FUCO, CARRA or NaOAc was added. The cell culture dish was covered and kept at RT for up to 30 days. Fig. 12 shows a photograph of the experiment involving **(2)** after six days of reaction.

**Fig. 12:**
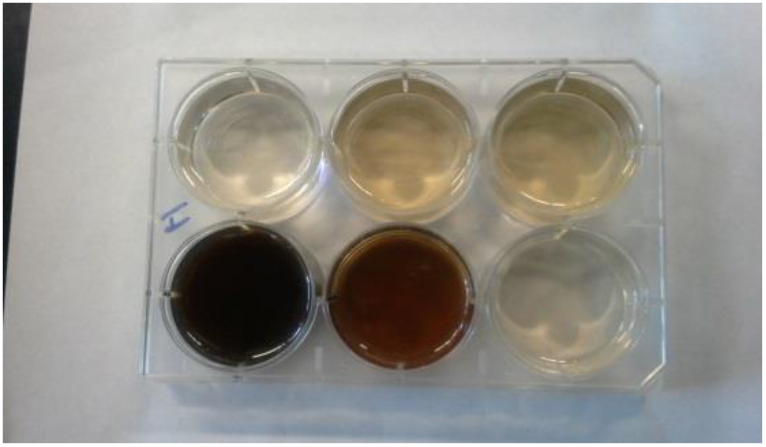
Photograph of reaction mixtures involving **(2)** and, top, from left to right, water, CS A and CS C or, bottom, left to right, FUCO, CARRA and NaOAc after six days at RT.

Dark colors had developed in the mixtures containing FUCO or CARRA and much lighter colors in all the other mixtures. Fig. 13 shows a photograph of the experiment involving **(3)** after thirty days of reaction.

**Fig. 13:**
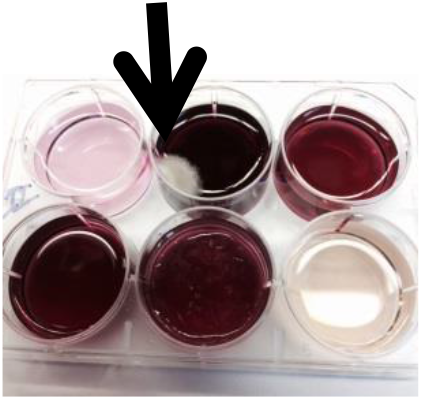
Photograph of reaction mixtures involving **(3)** and, top, from left to right, water, CS A and CS C or, bottom, left to right, FUCO, CARRA and NaOAc after thirty days at RT. The arrow points to mold growth that had occurred in the mixture containing CS A.

Reaction mixtures containing **(3)** very slowly developed a distinct reddish color and after thirty days of reaction intense colors were observed in the mixtures containing CS A, CS C, CARRA or FUCO, but not in the mixtures containing NaOAc or the control experiment. Mold growth had occurred in the mixture containing CS A (indicated with the arrow in Fig. 13), but not in any of the other mixtures. UV_Vis spectra of aliquots from select mixtures were recorded and are shown in Fig. 14.

**Fig. 14:**
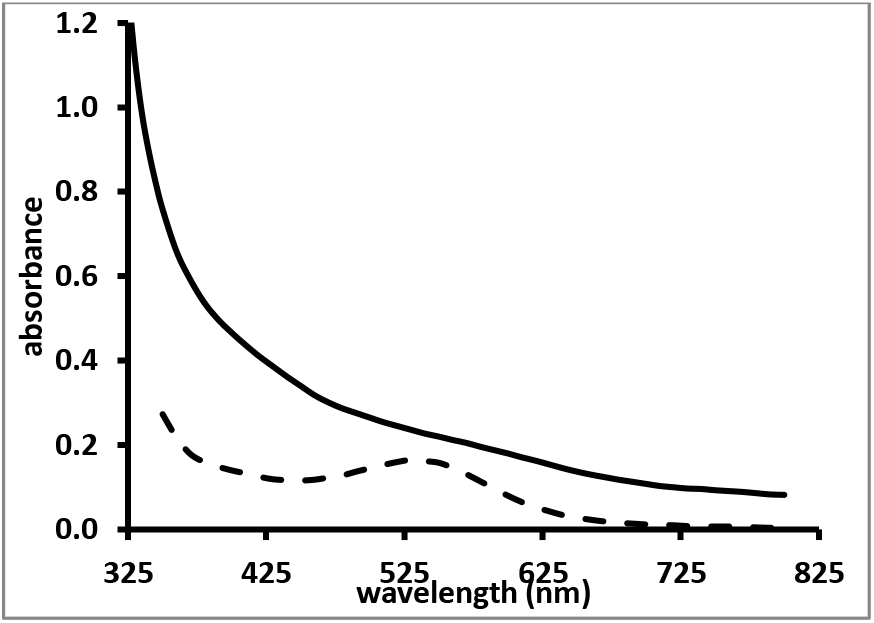
UV_Vis spectra of reaction mixtures involving **(2)** and FUCO (solid line) and **(3)** and CS C (dashed line) from the reaction mixtures shown in Fig. 12 and 13 respectively.

The spectrum of the reaction mixture involving **(2)** and FUCO showed enhanced absorbance in the entire visible range of the light spectrum, without any distinct features in the spectrum. Select reaction mixtures involving **(3)** yielded spectra with a distinct absorbance maximum around 530nm. Select reaction mixtures involving **(2)** were dialyzed against water and this purification was monitored using RP-HPLC analyses to ensure that all unreacted **(2)** had disappeared. The reaction mixture containing CS C and **(3)** was dialyzed against water, but all colored substances diffused out of the dialysis bag and after a couple of changes of water, the interior of the dialysis bag appeared to be colorless (results not shown).

It was presumed that the reaction between **(3)** and PS yielded a low molecular mass chromophore, but that no PS-pigment complex was generated. FT-IR spectra of select dialyzed and dried pigment materials made from **(2)** were recorded and typical results are presented in Fig. 15, panels A and B. Fig. 15, panel A, compares the FT-IR spectra of **(2)** and the pigment material made from FUCO and **(2)** in the absence of Cu^2+^. Fig. 15, panel B, compares the FT-IR spectra of FUCO and the same pigment material as in Fig. 15, panel A.

**Fig. 15:**
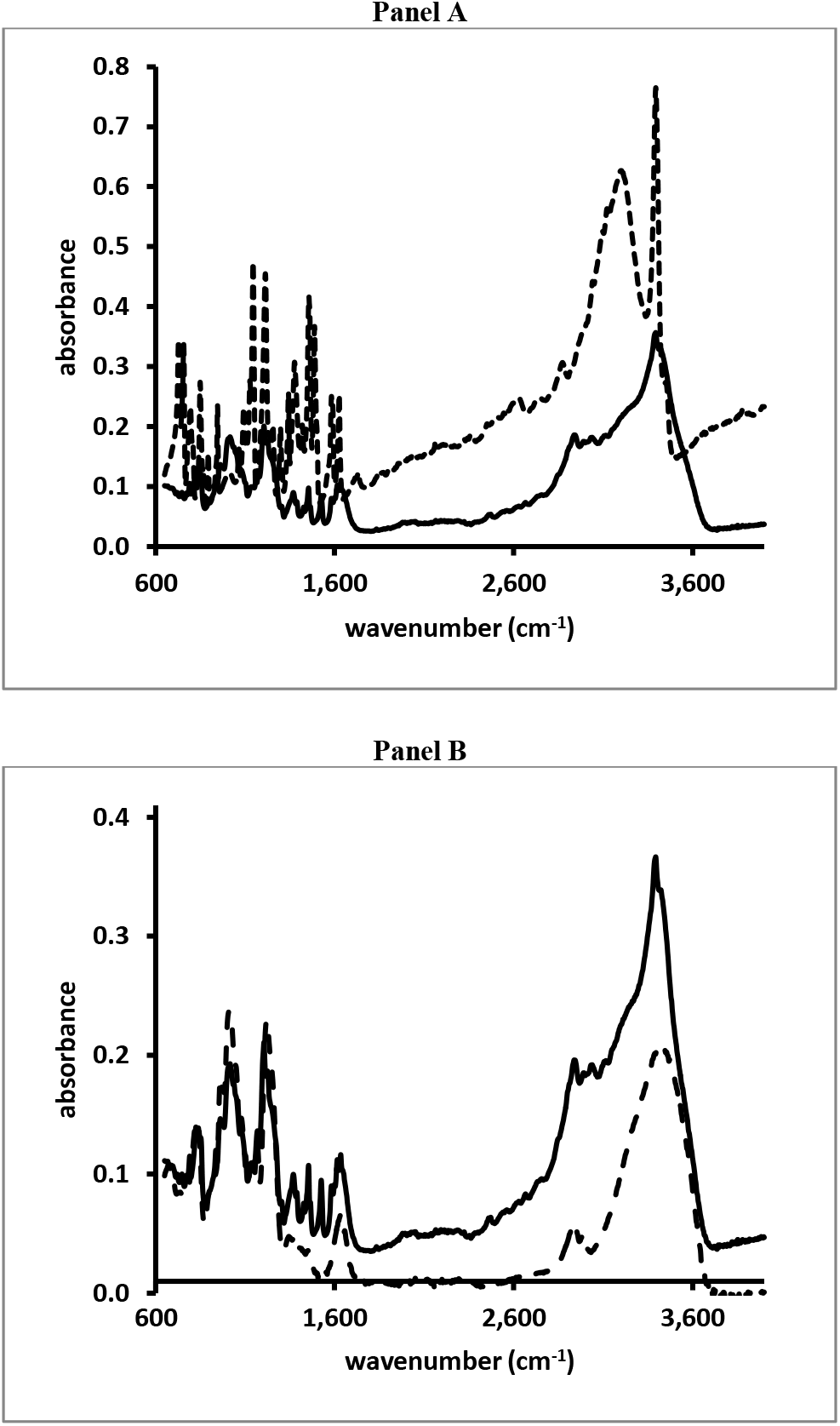
FT_IR scans of the pigment made from **(2)** and FUCO (solid line in both Panels) and **(2)** (dashed line in Panel A) or FUCO (dashed line in Panel B).

## 4. Discussion

As for CAs, select PS promote the auto-and Cu^2+^- mediated oxidation of **(1)** and related compounds resulting in the formation of pigment materials. The transformation of CAs into MN-like pigments is described through the so-called Raper-Mason scheme as discussed elsewhere.[14] In this scheme, CAs undergo oxidation and cyclization into indole types of structures prior to the formation of the MN pigment. Unlike the reactions involving CAs, the reactions involving **(1)** or **(2)** appeared not to generate a chromophore with distinct features in its UV_Vis spectrum prior to the generation of the darkly-colored pigment. Although, for both compounds, the oxidation reactions did lead to the formation of, as of yet unidentified, reaction products that could be resolved from **(1)** or **(2)** and other compounds through RP-HPLC or SEC analyses. The oxidation of **(3)** in the presence of select PS did lead to the formation of a chromophore with distinct features in its UV_Vis spectrum (see Fig.13 and 14), but this chromophore appeared not to react further to a high molecular mass type of pigment as CAs and **(1)** or **(2)** do.

The same types of PS that are capable of promoting the transformation of CAs into MN-like pigments, appear to enhance the oxidation of **(1)** and **(2)** into pigmented substances that, in color and SEC profile, resemble the MN-like pigments generated from CAs. Although autooxidation of both compounds, particularly in the presence of CARRA or FUCO, can be achieved, the reactivity is greatly enhanced with increasing concentrations of Cu^2+^ (see Fig.3 and 4).

SEC analysis of select pigmented materials generated from **(1)** in the presence of PS indicate that a fraction of the pigment is associated with the PS material and that another fraction of the pigment is associated with some material with much higher molecular mass than the original PS (see Fig. 9). MNs formed from CAs have been observed to exist in nanoparticulate form.[15, 16] Thus, it is not inconceivable that pigments generated from **(1)** in the presence of PS could exist as PS-stabilized nanoparticles.

The Cu^2+^-mediated oxidation of **(1)** in the presence of PS like CS A leads to the formation of an unidentified reaction product that can be resolved from **(1)** and PS materials through SEC analysis (see Fig. 5 and 9). Jones et al. and Dai et al., in their studies of the Cu-induced oxidation of **(1)**, identified a dimeric compound, 5,5’- dihydroxy-4,4’-bitryptamine or DHBT and its isomers, as reaction products.[12, 17] However, the unidentified reaction product generated in our experiments is not removed through dialysis using a membrane with MWCO of 3.5kDa (see Fig. 9), hinting that it represents a high molecular mass compound. This suggests that the unknown product generated in our experiments is unlikely to be DHBT.

The FT-IR spectra obtained of pigmented materials generated from **(1)** (see Fig. 10) or from **(2)** (see Fig. 15) in the presence of select PS, contains the presence of a distinct peak that is not present in the FT-IR spectra of the PS and that is present in the spectra of **(1)** or **(2)**. Comparing the spectrum of CS A to the spectrum of the CS A/pigment material in Fig. 10, panel B, the sharp peak at about 3,390 cm^-1^ stands out. Such a sharp peak is also present in the spectrum of **(1)** as seen in Fig. 10, panel A. Comparing the spectrum of FUCO to the spectrum of the FUCO/pigment material in Fig. 15, panel B, a small, sharp peak at about 3,393 cm^-1^ can be observed. Such a sharp peak at that wavenumber is present in the spectrum of **(2)** as shown in Fig. 15, panel A. FT-IR spectra of indole types of compounds show sharp peaks around 3,390 cm^-1^ and these have been attributed to an asymmetric NH stretching band.[18, 19] Thus, FT-IR spectra of the pigments generated from **(1)** or **(2)** do contain chemical signatures different from those of the PS and these cannot be attributed to **(1)** or **(2)** as any unreacted starting material was removed through the dialysis process.

Apart from its importance in neurochemistry, the links between **(1)**, its receptors and transporters and the immune system is the focus of intense research as reviewed elsewhere.[20] Peripheral sources release **(1)** into the bloodstream or lymphatic tissues for interaction with the various components of the innate or adaptive immune system.[20] The enterochromaffin (EC) cells of the gastrointestinal tract (GI) are such an important peripheral source of **(1)**.[21] It is interesting to note that the presence of a greyish-brown pigment in the cytoplasm of such cells had been described more than forty years ago.[22] Among the multiple effects of **(1)** on the components of the immune system, is its capacity to modulate the release or synthesis of pro-inflammatory cytokines like IL-1β.[20]

In addition, **(1)** is capable of promoting the release of IL-6, e.g., from rat adrenal zona glomerulosa cells.[23] Therefore, we briefly investigated whether pigments derived from **(1)** could affect the release of IL-1β or IL-6 from immune cells. As shown in Fig. 11, panel A, pigment material generated from **(1)** in the presence of CS A significantly increased the amount of IL-1β from immune cells in a dose-dependent fashion. CS A had only a modest effect on the release of IL-1β. In contrast, CS A promoted the release of IL-6 in a dose-dependent fashion. The pigment material generated from **(1)** in the presence of CS A, tested at the lowest concentrations, appeared to have a similar effect on the IL-6 release as CS A, while when tested at higher concentrations, the IL-6 release in the presence of the pigment material was reduced with about 30% compared to CS A.

## Acknowledgements

This research and Astiney Clark were in part supported by a grant from the US Department of Education [#P031B090214]. Part of the research was supported by grants U54CA163066 and 2T34GM007663 from the National Institutes of Health. Noor Alatas was supported by the Saudi Arabian Cultural Mission to the USA.

